# Regulation of Cardiomyocyte Adhesion and Mechanosignalling Through Distinct Nanoscale Behaviour of Integrin Ligands Mimicking Healthy or Fibrotic ECM

**DOI:** 10.1101/2022.03.13.483961

**Authors:** William Hawkes, Emilie Marhuenda, Paul Reynolds, Caoimhe O’Neill, Pragati Pandey, Darren Graham Samuel Wilson, Mark Freeley, Da Huang, Junquiang Hu, Sasha Gondarenko, James Hone, Nikolaj Gadegaard, Matteo Palma, Thomas Iskratsch

## Abstract

The stiffness of the cardiovascular environment changes during ageing and in disease and contributes to disease incidence and progression. Changing collagen expression and crosslinking regulate the rigidity of the cardiac ECM. Additionally, basal lamina glycoproteins, especially laminin and fibronectin regulate cardiomyocyte adhesion formation, mechanics and mechano-signalling. Laminin is abundant in the healthy heart, but fibronectin is increasingly expressed in the fibrotic heart. ECM receptors are co-regulated with the changing ECM. Due to differences in integrin dynamics, clustering, and downstream adhesion formation this is expected to ultimately influence cardiomyocyte mechanosignalling; however details remain elusive. Here we sought to investigate how different cardiomyocyte ligand/integrin combinations are affecting adhesion formation, traction forces and mechanosignalling, using a combination of uniformly coated surfaces with defined stiffness, PDMS nanopillars, micropatterning and specifically designed bionanoarrays for precise ligand presentation. Thereby we find that neonatal rat cardiomyocytes (which express both laminin and fibronectin binding integrins) adhesion nanoscale organisation, signalling and traction force generation are strongly dependent on the integrin/ligand combination. Together our data indicates that the presence of fibronectin in combination with the enhanced stiffness in fibrotic areas will strongly impact on the cardiomyocyte behaviour and influence disease progression.

## Introduction

The cardiac extracellular matrix (ECM) provides structural support for cardiomyocytes, the contractile cells of the heart. Additionally, the chemical and mechanical composition of the ECM is sensed by the cardiomyocytes and influences their differentiation, maturity, and function. Importantly, the ECM composition, structure and mechanics change during development and in heart disease. Changes include different collagen isoforms, collagen crosslinking, upregulation of fibronectin in the basal lamina and increasing myocardial stiffness from ∼1 kPa in the foetal heart to ∼ 10kPa in the healthy adult heart and 50-150kPa in the diseased, fibrotic myocardium[1]. These changes are sensed through different pathways, but especially through integrin adhesion and downstream mechanosignalling [2-5]. The transmembrane integrin receptors enable inside-out and outside-in mechanotransduction[6]. On the outside of the cell, they bind to extracellular matrix molecules, such as laminin, fibronectin, or collagen, depending on the integrin isoforms. On the cytoplasmic face they connect to the actin cytoskeleton via an adhesion complex with talin at its core. Actomyosin forces on talin can open cryptic binding sites for vinculin (depending on the stiffness of the ECM), which reinforces the adhesion through providing further connections to the actin cytoskeleton[6, 7].

Cardiomyocyte integrins are primarily located at specific circumferential adhesion sites, the so called costameres, although in 2D culture integrins are further found in focal adhesion like structures[8]. The cardiomyocyte integrin expression profile is highly regulated, and specific expression patterns are activated during developmental, neonatal, healthy adult and diseased states [8-13]. Specifically, the laminin binding α7β1 integrin becomes the primary receptor in adulthood however, the fibronectin binding α5β1 integrins are the predominant subtype during development and in infarcted, or ischemic hearts, where increasing amounts of fibronectin are deposited by cardiac fibroblasts [1, 13].

However, different integrins vary in their ligand binding characteristics, dynamics, and nanoscale organisation[1, 14-16] and therefore, not only ECM stiffness but also the specifics of the ECM ligand/integrin are expected to influence the adhesion mechanosignalling. Indeed, changes to adhesion composition have been linked to postnatal heart development as well as a maladaptive response and progression to heart failure, but details remain unclear[1]. To address this, we sought here to investigate how different cardiomyocyte ligand/integrin combinations are affecting adhesion formation, traction forces and mechanosignalling, using a combination of: i) uniformly coated surfaces with defined stiffness, ii) PDMS nanopillars, iii) micropatterning, and iv) specifically designed bionanoarrays for precise ligand presentation.

Using this combined approach, our results consistently demonstrate that NRCs on fibronectin have enhanced spreading than cells cultured on laminin, at physiological and fibrotic stiffness. Moreover, NRCs cultured on fibronectin exhibit greater traction forces than cells cultured on nanopillars coated with laminin. Additionally, micropatterning revealed greater vinculin enrichment to fibronectin adhesions, suggesting altered rigidity sensing and mechanotransduction. Finally, our use of DNA origami bionanoarrays identifies unique properties of both laminin or fibronectin (RGD) binding integrins, especially differences in minimal local and global ligand concentrations needed for stable adhesion formation and spreading. Together our results demonstrate that integrin-ligand paring critically influence the mechanosignalling of cardiomyocytes.

## Results

### Cardiomyocyte morphology and spread area are affected by the extracellular matrix composition

Because of changing extracellular matrix (ECM) composition and co-regulation of integrin isoforms we first wanted to establish differences in cardiomyocyte phenotype dependent on the ECM stiffness and composition. Neonatal cardiomyocytes (NRC) are a popular model for cardiomyocyte studies due to their relatively maturity (e.g. compared to standard IPSC-CM models) and ease of culture (e.g. compared to adult cardiomyocytes which start to dedifferentiate immediately after plating[17]). Indeed, NRC are well suited to study the interaction of cardiomyocytes with different extracellular matrix components since they express high levels of both laminin and fibronectin binding integrins (Supplementary Figure S1)[12].

We first analysed cell area and shape of NRC when plated on PDMS surfaces with different stiffness that were coated with either fibronectin or laminin (Fig 1). On fibronectin cells started showing a characteristic mechanosensitive behaviour after 48h. In agreement with our previous findings, largest cardiomyocyte areas and strongest α-actinin staining were found on fibronectin on 20kPa surfaces[5], whereas cell shape changes were more pronounced on 130kPa with an increased solidity and form factor, in agreement with a more spindle-shaped appearance (Fig 1D,E). In contrast, cardiomyocytes on laminin did not show significant stiffness-dependent differences after 48h. However, after 72h cardiomyocytes displayed a similar mechanosensitive response also on laminin, with largest areas and strongest actinin staining on 20kPa and progressively increasing solidity and form factor with increasing stiffness (Fig 1).

**Figure 1.**
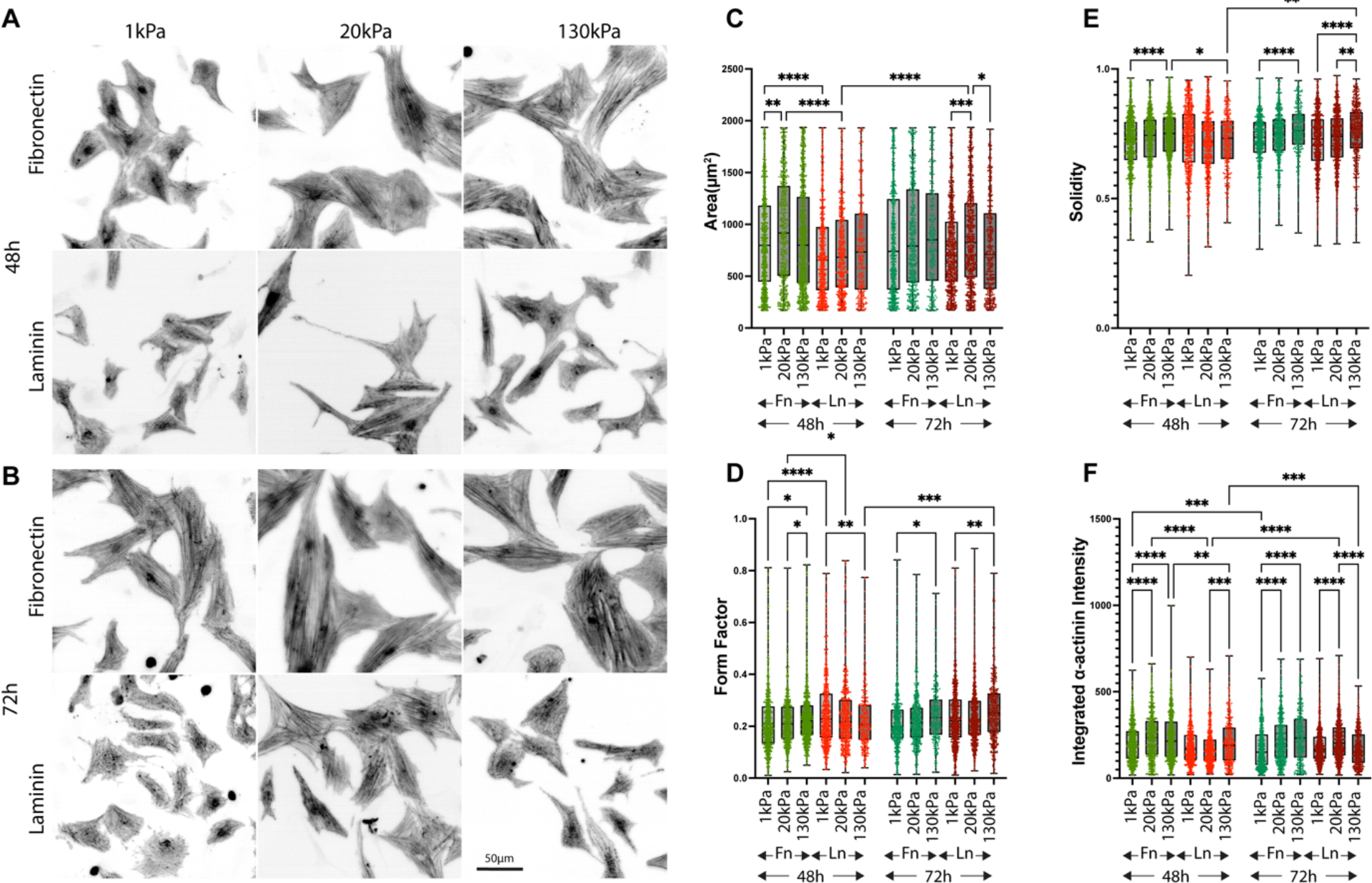
NRC spreading on PDMS substrates coated with fibronectin (A) or laminin (B). Quantified Cell Area after 48 and 72 hours in culture. n = 3 independent experiments, > 400 cells per condition. * = p < 0.05, ** p < 0.01, *** = p < 0.001, *** = p < 0.0001. P values from one-way ANOVA (fibronectin vs laminin for each condition) and Tukey correction for multiple comparisons. Scale bars = 50 *μ*m. D-F) Cytoskeletal morphology of NRCs cultured on fibronectin of laminin for 24 hours. Cells on fibronectin (A) and laminin (B) coated substrates were stained for α-actinin and cell morphology (C-E) and α-actinin intensity (F) were quantified from three independent experiments with > 300 cells per condition.

### Cardiomyocyte adhesion forces are increased on fibronectin compared to laminin

The different temporal evolution of mechanosensitive behaviour suggested differences in adhesion formation and signalling depending on the ECM ligand. Since adhesion formation and cell morphology are closely related to adhesive forces, we next investigated cardiomyocyte contraction forces that were transmitted through the different integrin/ECM combinations, using nanopillars (Fig 2). Movements of pillars, coated with either fibronectin or laminin were recorded at frame rates of >10fps. From the pillar movements, the maximum displacement (systole) was compared to the subsequent minimum (diastole) and to the non-displaced pillars and the contraction force was calculated as difference between systolic and diastolic force (Fig 2A-C). Consistent with a difference in spread area and mechanosensing, we found indeed higher forces (both systolic and diastolic) from the NRCs that were plated on the fibronectin coated pillars.

**Figure 2.**
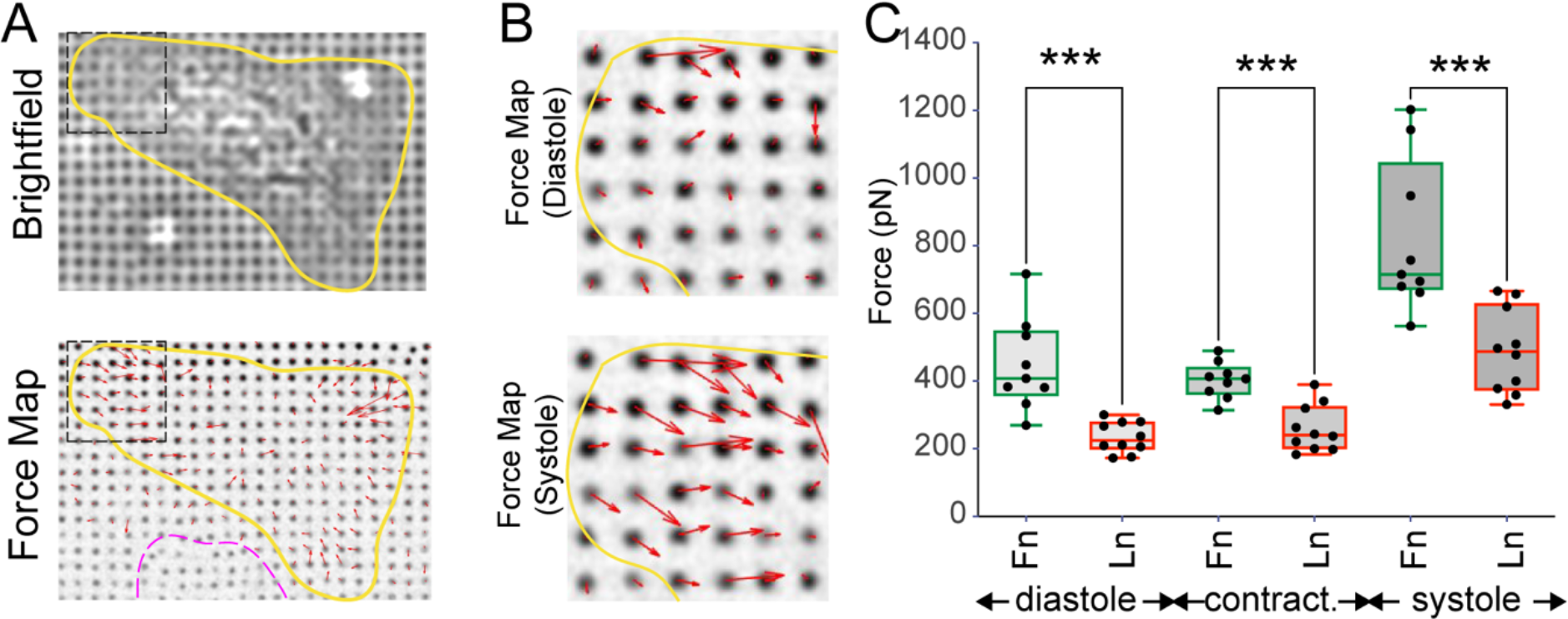
NRC generate higher traction forces on fibronectin vs laminin. A-C) NRC traction forces were analysed using PDMS nanopillars coated with quantum dots. A) Bright field images were taken prior to fluorescent movies to enable identification of cell area, while quantum dot movies enabled the calculation of the force maps. Yellow indicates cardiomyocyte; magenta indicates a fast-migrating cell as detected by deformation of different pillars on different frames. Only cardiomyocyte forces were quantified. B) Zoom into box from (A) for detailed force maps during diastole and systole. C) Analysis of diastolic and systolic pillar displacements after 48 hours in culture. n = 10 & 9 cells for laminin and fibronectin respectively from 3 independent experiments. *** p < 0.001. P values from students t-test (fibronectin vs laminin).

### Micropatterning indicates enhanced adhesion formation on fibronectin vs laminin

Micropatterning perpendicular lines of different proteins offers an elegant way to analysis the differential interaction of cells with two or more receptor ligands, which we previously employed to study the immune synapse formation in T-cells[18]. This method allows to directly compare the adhesive strength, by analysing the relative alignment of the cells with one ligand over the other. Additionally, it allows the measurement of enrichment of adhesion proteins over the ligand lines. Therefore, we decided here to pattern perpendicular lines of fibronectin versus laminin (Fig 3). To further investigate the mechanosensing we decided to pattern on PDMS of different stiffness. Confirming the results from the uniformly coated surfaces we found stronger alignment of the NRC with the fibronectin lines compared to the laminin lines. This result was more pronounced on 130kPa vs 20 kPa (Fig 3B).

**Figure 3.**
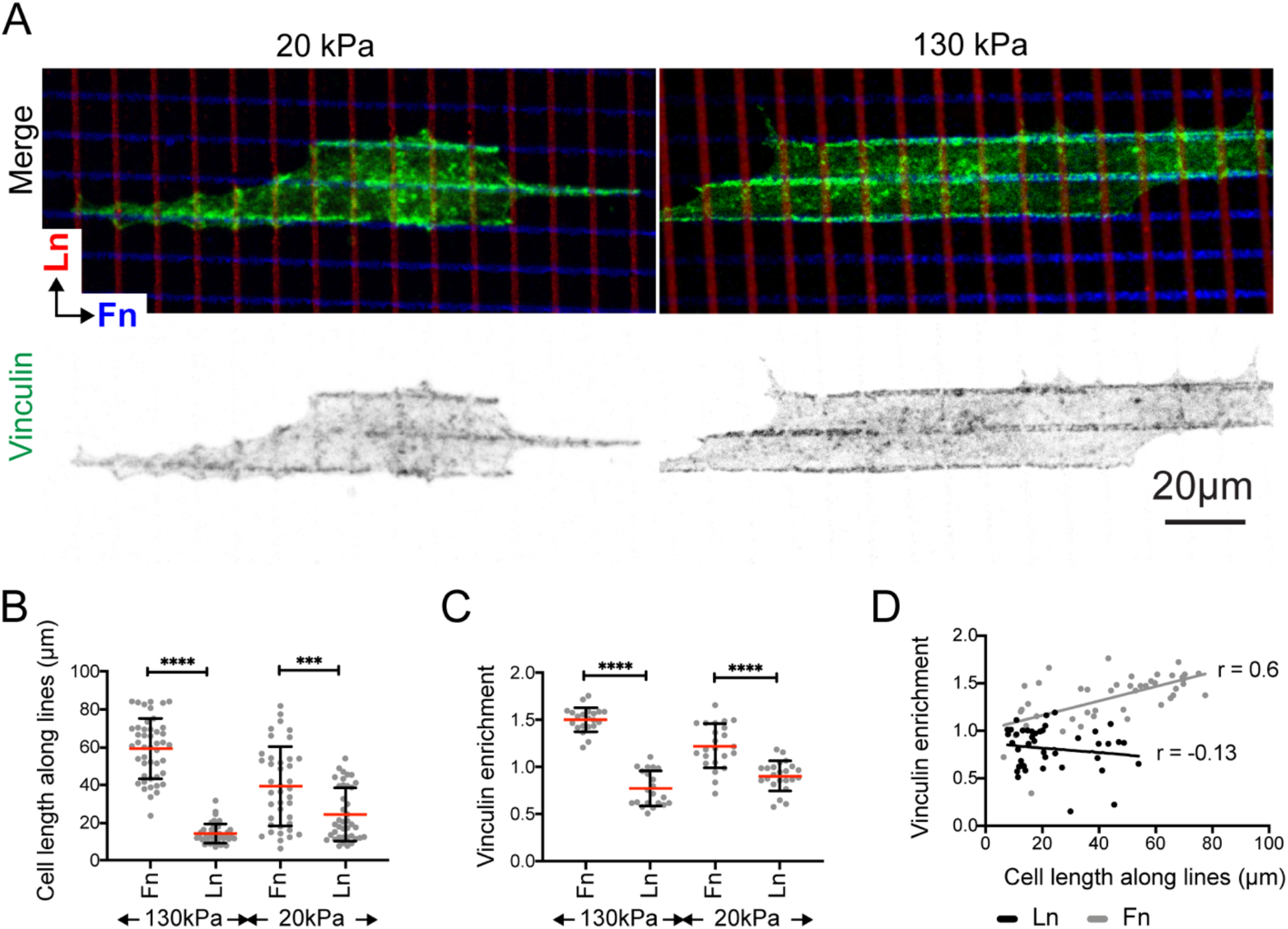
Vinculin enrichment to fibronectin indicates adhesion reinforcement. D) Representative images E) Quantification of cell extent along the respective directions of the grids. n = 46 and 40 cells for 130 kPa and 20 kPa conditions. F) Analysis of vinculin enrichment to the grids and G) Persons correlation coefficient of vinculin enrichment vs cell extent. n = 21 & 23 cells at 130 kPa and 20 kPa conditions. **** = p < 0.0001. P values from students t-test.

Forces on talin reveal cryptic binding sites for vinculin and adhesion re-inforcement[19]. Therefore, the respective amount of vinculin can be used as a surrogate measurement of adhesion reinforcement. Indeed, vinculin showed a strong enrichment along the fibronectin lines that was further enhanced at 130kPa vs 20kPa (Fig 3C). Moreover, we found a correlation between vinculin enrichment and spread length along the fibronectin lines, but not between vinculin enrichment and spread length along the laminin lines (Fig 3D). Together these data indicated that adhesion strength and adhesion forces are more pronounced on fibronectin compared to laminin and result in enhanced vinculin recruitment and adhesion reinforcement.

### DNA Origami for controlled presentation of receptor ligands

Since our findings so far demonstrated unique adhesive behaviours of NRCs when cultured on fibronectin compared to laminin, we hypothesised that these might be related to the specific adhesion structure and composition. Indeed, investigations have shown key differences in activation [20] and clustering [15] of different integrin subtypes.

To understand the role of these factors for the regulation of cardiomyocyte adhesion to the ECM, we decided to employ DNA origami as a tool to control the distance of receptor-ligands with nanometre resolution (Fig 4-6). To investigate the nanoscale organisation and clustering of fibronectin and laminin binding integrins in cardiomyocytes, we used DNA origami presenting 0, 6, or 18 RGD (for fibronectin), or IKVAV peptides (to mimic laminin). At a side length of 127nm per DNA origami triangle, 6 ligands corresponded to an inter-ligand spacing of approximately 60nm, 12 ligands to ∼30nm inter-ligand spacing, and 18 ligand were spaced by ∼20nm each.

**Figure 4:**
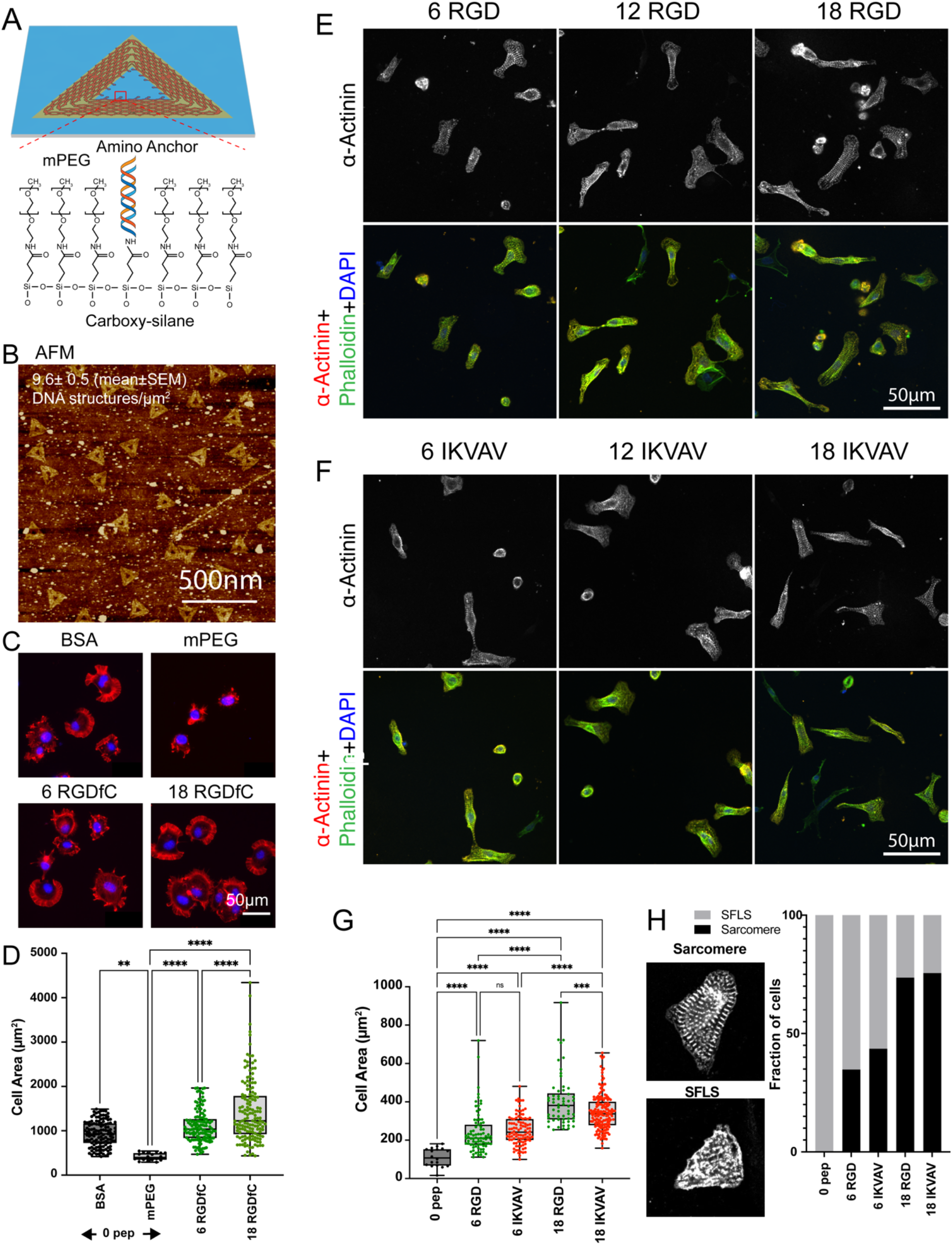
Spreading on random Origami scales with integrin ligand spacing. A-D) Optimisation of conditions using mouse embryonic fibroblasts cultured on random DNA origami arrays. A) mPEG-amine was used to block remaining carboxyl groups after attachment of DNA origami via amino anchors to carboxyl-terminated silanes. B) AFM image of randomly positioned DNA origami. C) mPEG-amine blocking prevent non-specific adhesion to the background on DNA origami functionalised with 0, 6, or 18 RGD. Red: Phalloidin, Blue: DAPI. D) Quantification of n=3 independent experiments. One-way ANOVA with Tukey correction for multiple comparisons, ** = p<0.01, **** = p<0.0001. E-H) NRCs were cultured on RGD or IKVAV functionalised nanopatterns for 24 hours. E,F) Representative images of NRCs spreading on the RGD (E) and IKVAV (F). Spreading was quantified by measuring cell area (G). H) maturity was assessed, by counting the number of cells containing stress fibre like structures (SFLS) or additionally contained mature sarcomeres. **** = p<0.0001. Data pooled from 3 independent experiments.

DNA origami were cast on glass substrates at a concentration of 2 nM and crosslinked via amino-anchors to carboxyl-terminated silanes of the functionalised glass, leading to a density of 9.6 ± 0.5 DNA structures per *μ*m^2^ (Supplementary Figure S2). Remaining carboxyl-groups were subsequently blocked by crosslinking to mPEG-amine and successful passivation was confirmed by plating of mouse embryonic fibroblasts (MEFs, Fig 4A-D).

Like the MEFs, NRCs were unable to spread on zero peptide controls (with mPEG passivation), confirming that non-specific adhesion was minimal. Upon presentation of 6 peptides per origami (∼60 ligands per *μ*m^2^), NRCs started to spread on both RGD and IKVAV, with no significant differences observed between the two peptides. In general, NRCs were small on 6xRGD and 6xIKVAV arrays (240 ± 107 *μ*m^2^ and 252 ± 76 *μ*m^2^ for RGD and IKVAV respectively) and most NRC exhibited primarily stress-fibre like structures (SFLS) with few mature sarcomeres (Fig 4E-H).

In contrast, the cell area was considerably larger on both RGD and IKVAV arrays presenting 18 ligands (∼180 ligands per *μ*m^2^). Here, significantly more cell spreading was observed on RGD compared to IKVAV. Nevertheless, most cells on both 18xRGD and 18xIKVAV were able to form mature sarcomeres. These findings clearly demonstrate that both RGD and IKVAV functionalised DNA origami facilitate NRC spreading and cytoskeletal maturation in a dose dependent manner, whereby presentation of RGD peptides resulted in larger cell areas compared to IKVAV, but comparable maturity.

### Different nanoscale behaviour of integrins regulate cardiomyocyte adhesion

The investigation of NRCs on randomly positioned DNA origami functionalised with RGD or IKVAV demonstrated that this platform can mediate NRC spreading and adhesion. However, the random arrays can only provide information on integrin clustering relative to the area of a single origami structure (∼85 nm^2^) with little control over the global positioning of the origami. Nanopatterned DNA origami (here bionanoarrays) overcomes these limitations by providing precise control of both global and local integrin-ligand placement. Specifically, we generated circular pattern with 150nm diameter, using e-beam lithography. After sialinisation (using carboxyl-terminated silanes as above) and removal of the remaining resist, DNA origami were conjugated to the carboxyl groups, present only at the specified locations (Fig 5A, AFM images).

**Figure 5.**
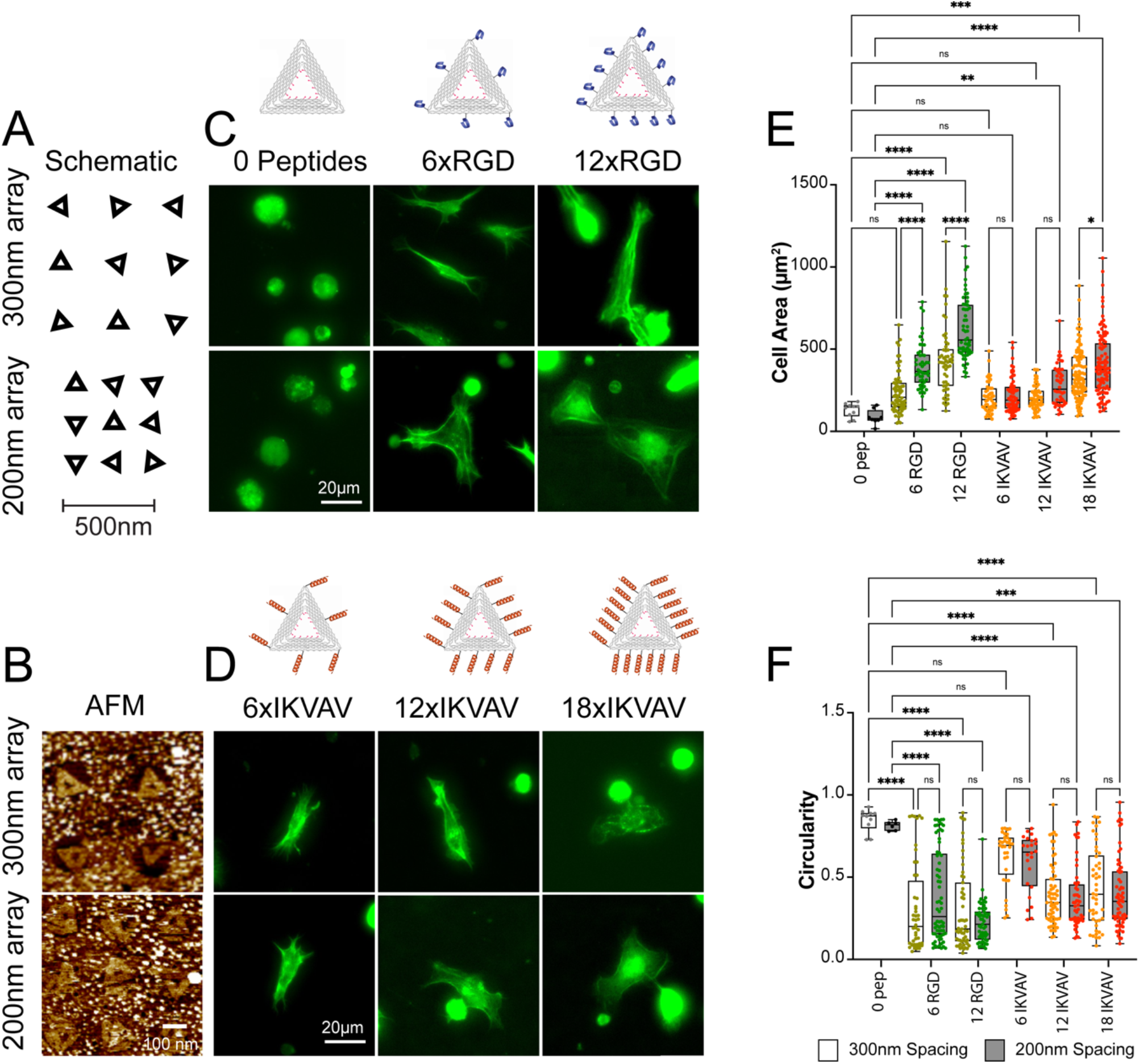
Bionanoarrays for study of nanoscale adhesion organisation. A) schematic of bionanoarrays set up; since circular patterns are used, distance but not orientation of the origami triangles can be controlled B) AFM images of DNA origami bionanoarrays C,D) Representative images of NRCs spreading on the RGD (C) and IKVAV (D) nanopatterns. The schematic on top shows the approximate placing of the peptides. Cells were stained with phalloidin and analysed for cell area (E) and circularity (F). Differences between peptide numbers and origami spacing were evaluated with a Two-way ANOVA with Tukey corrections for multiple comparisons, ** = *p*<0.01, *** = *p*<0.001, **** = *p*<0.0001. Data from 3 independent experiments. N = >50 cells per condition.

To compare the global and local minimal ligand densities and clustering behaviours of different cardiomyocyte integrins, bionanoarrays were fabricated with 0 peptides, 6, or 12 RGD or 6, 12 or 18 IKVAV peptides in arrays with inter-origami spacings of 200 nm and 300 nm (Fig 5; supplementary table 1 summarises the local and global ligand properties of each nanopattern configuration). Bionanoarrays with zero peptides were employed as negative controls.

Using such bionanoarrays enabled us to specifically study minimal inter-ligand spacings and global densities in order to facilitate cardiomyocyte spreading. Again, on zero peptide controls, few cells adhered to the substrates, suggesting a low level of non-specific adhesion. The addition of 6xRGD peptides (∼60 nm spacing) resulted in an increase in cell attachment and cell area and a decrease in circularity (Fig 4). This indicated that the local ligand density was sufficient to allow at least some level of spreading (similar to previous work in migratory cells)[21].

Nevertheless, on arrays with 200nm cluster spacing (6xRGD/200nm; 150 peptides/*μ*m^2^), the cardiomyocyte area was significantly larger than on 6xRGD/300nm bionanoarrays (88 peptides/*μ*m^2^). Increasing the number of RGD peptides to 12 (decreasing inter-ligand spacing to ∼30 nm) resulted in a further increase in cell area on both, 12xRGD/200nm and 12xRGD/300nm arrays. Interestingly, NRC area on the 12xRGD/300nm bionanoarray was comparable to the 6xRGD/200nm array. Since both arrays had comparable overall ligand densities (Supplementary Figure S4, 176 peptides/*μ*m^2^ vs 150 peptides/*μ*m^2^), this result clearly suggested a dominance of global over local ligand density for fibronectin binding integrins in cardiomyocytes.

On IKVAV bionanoarrays in contrast, only few cells were spread on either 6xIKVAV/200nm or 6xIKVAV/300nm arrays, suggesting insufficient integrin clustering and adhesion formation. This was also apparent from the cell morphology with higher circularity indicating overall lesser amount of spreading and maturation compared to the corresponding RGD arrays (Figure 5D). With 12xIKVAV peptides, a significant cell area increase was only observed on 12xIKVAV/200nm arrays, although the change in cell shape was testament for an activation of integrin signalling and adhesion stabilisation also on 12xIKVAV/300nm arrays (Fig 5C,D). Together, this suggested that cardiomyocyte laminin binding integrins required a smaller inter-ligand distance of ∼30nm vs ∼60nm to enable integrin adhesion formation and spreading.

Further spreading was facilitated by adding 18xIKVAV ligands per DNA origami (∼20nm inter-ligand spacing). Here, cardiomyocytes displayed again larger cell areas on 18xIKVAV/200nm arrays compared to 18xIKVAV/300nm arrays. However, spread area was still significantly less than on 12xRGD/200nm arrays (and comparable to 12xRGD/300nm arrays). Importantly, cardiomyocyte area was larger on 18xIKVAV/300nm compared to 12xIKVAV/200nm bionanoarrays, even though the former had a lower global ligand density of 208 vs 300 peptides/*μ*m^2^. This was in contrast to RGD bionanoarrays and suggested a reduced role of global compared to local densities for laminin vs fibronectin binding integrins in neonatal cardiomyocytes.

In summary the bionanoarrays indicated different minimal inter-ligand spacings (30nm vs 60nm for IKVAV and RGD respectively) and different contributions of local and global ligand densities that are needed for activating integrin signalling and supporting stable cell adhesion and spreading.

### The integrin ligand affinity is balanced with the cytoskeletal contractility

Since we found differences in spread area, cytoskeletal arrangements and adhesion signalling, we next thought to investigate the source of this discrepancy. Previously we found a role for myofibrillar and non-myofibrillar tension in regulating cardiomyocyte adhesion formation and mechanosensing [5]. To investigate this further, NRCs spreading on random 18xRGD or 18xIKVAV arrays were treated with the contractility inhibitors blebbistatin (to inhibit myosin II) and Y-27632 (to inhibit ROCK) or contractility enhancers Calyculin-A (inhibits myosin-light-chain phosphatase from dephosphorylating myosin[22]) and Omecamtiv Mercabil (OM, binds to the catalytic domain of myosin and increases cardiac contractility[23])(Figure 6).

**Figure 6.**
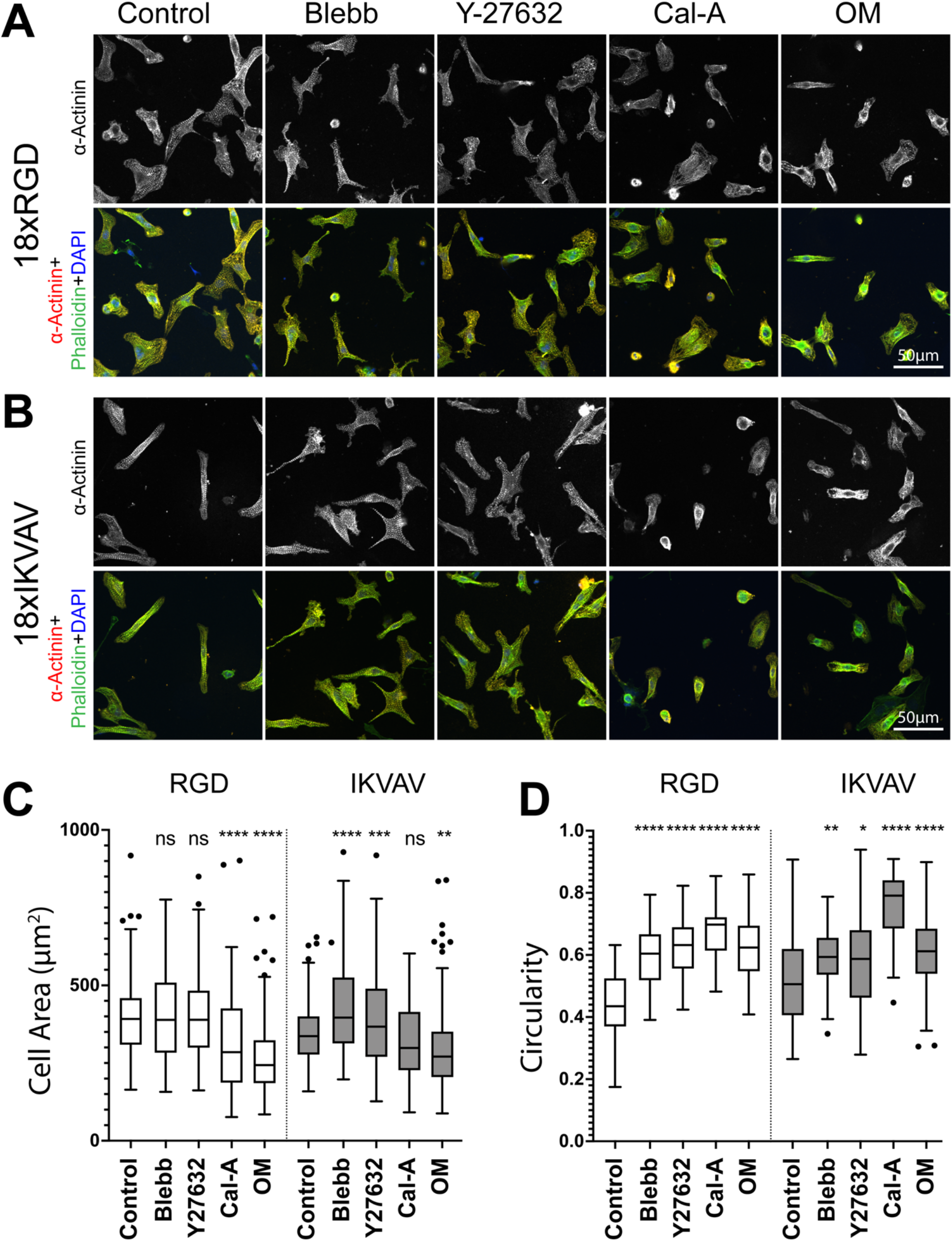
Effect of cytoskeletal contractility inhibitors and activators on NRC spreading on DNA origami. NRCs were cultured on random origami arrays presenting 18xRGD or 18xIKVAV peptides and treated with drugs to inhibit (Blebbistatin (5 *μ*m), Y-27632 (10 *μ*m)) or promote cytoskeletal contractility (Calyculin-A (100 nM), Omecamtiv Mercabil – OM (500 nM). A) Representative images of NRCs immunostained for α-actinin and phalloidin. B) Quantification of cell area. C) Quantification of circularity. One-way ANOVA with Tukey correction for multiple comparison, * = *p*<0.05, *=*p*<0.01, ***=*p*<0.001, ****=*p*<0.0001. N= 3 independent experiments.

Reducing contractility with blebbistatin or Y-27632 resulted in a significant increase in cell area on IKVAV (but not on RGD). This result suggests that the adhesions to IKVAV at the specific ligand concentrations employed here, are unstable at the existing levels of contractility. Therefore, reducing the contractility stabilised adhesions and facilitated spreading. In contrast, drugs promoting contractility resulted in a decrease in cell area on both RGD and on IKVAV functionalised DNA origami nanoarrays. The decrease was stronger with OM and only this treatment showed a significant effect on IKVAV modified surfaces.

Together, these results suggest that cytoskeletal contractility, and integrin ligand affinity need to be well balanced to support cell spreading. IKVAV/laminin-binding integrins overall require higher density of ligands to support adhesion formation and downstream signalling and hence cannot support the contractility of neonatal cardiomyocytes to the same extent as α5β1 integrin adhesions to RGD (see model in Fig 7).

**Figure 7.**
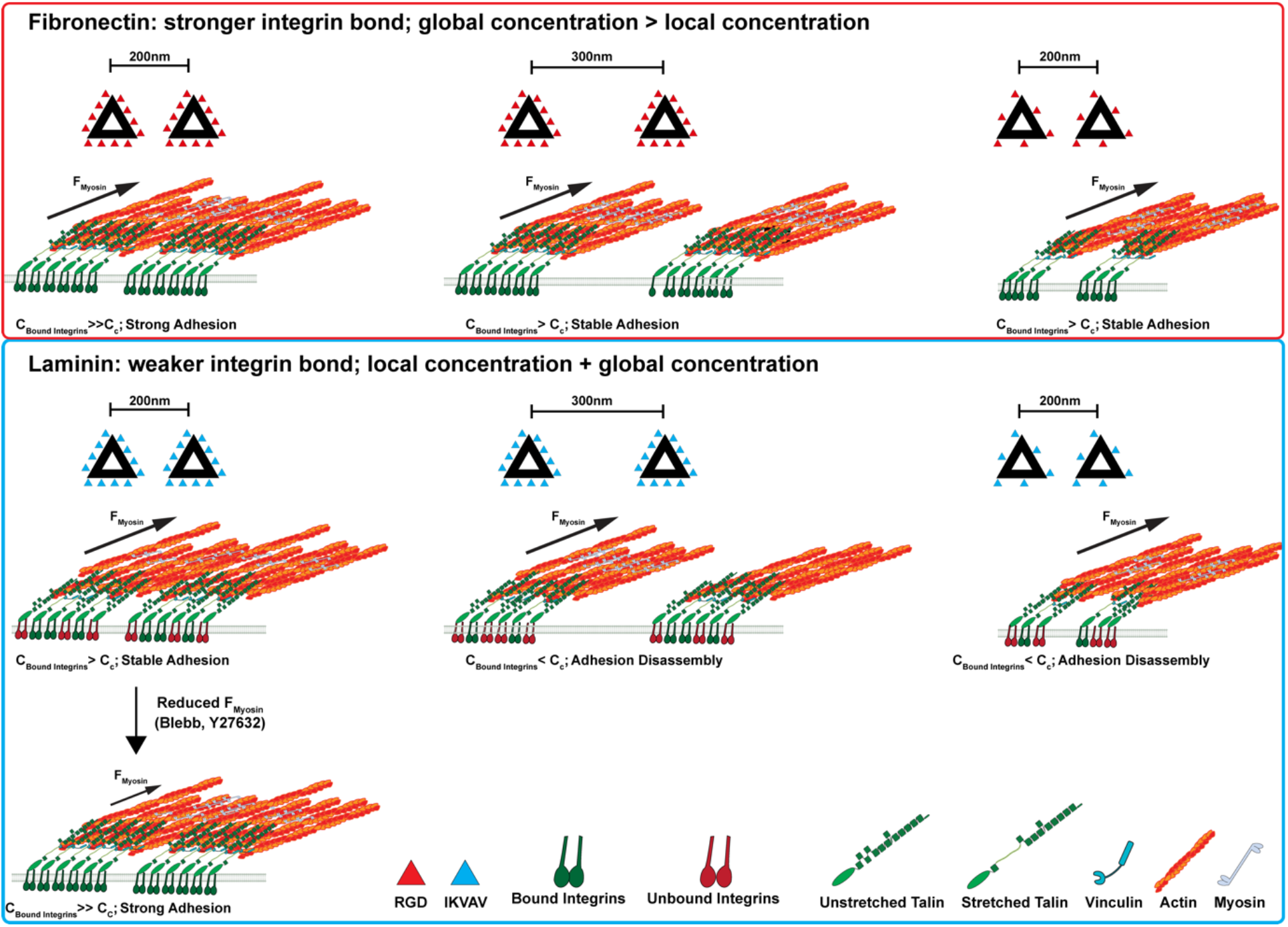
Model for nanoscale requirements of fibronectin and laminin binding integrins in cardiomyocytes. **Top panel:** fibronectin binding integrins establish relatively strong bonds to their ECM ligands and/or talin and cytoskeletal forces lead to talin stretching, vinculin binding and adhesion reinforcement (i.e. the concentration of bound integrins, C_Bound Integrins_, is larger than the required critical concentration, C_c_). Stable adhesion clusters can be formed with a smaller number of ligands and because of the higher stability of bonds, larger distances between clusters can be sustained. Overall, the global concentration is more important for determining the adhesion stability than the local concentration. **Bottom panel:** weaker bonds of laminin binding integrins either to the ECM ligand and/or talin result in more frequent bond ruptures. This results in an overall lower number of force-bearing integrin-talin links at a certain time (i.e. the concentration of bound integrins, C_Bound Integrins_, is small than the required critical concentration, C_c_). A larger number of ligands is required to stabilize the adhesion clusters. Hence local, as well as global concentrations of integrins determine the adhesion strength. Reducing the myosin forces through treatment with blebbistatin or Y27632 increases the bond lifetimes and adhesion stability.

## Discussion

During development and in disease, cardiomyocyte express fibronectin binding integrins (especially α5β1), while laminin binding integrins are the dominant subtypes in healthy adult cardiomyocytes [1, 10, 12]. Previous work demonstrated that different integrin subtypes have unique ligand binding and rigidity sensing properties, but the consequences integrin signalling, and cardiomyocyte phenotype are not well understood [1, 14-16, 24, 25]. The current investigation made use of an interdisciplinary approach to conduct a comparative analysis of neonatal rat cardiomyocyte spreading and integrin clustering on fibronectin and laminin functionalised substrates. We demonstrated that cardiomyocytes cultured on fibronectin exhibit stronger spreading, contractility and vinculin enrichment compared to cells cultured on laminin. We hypothesised that these differences could arise due to differential integrin clustering. Indeed, by implementing a single molecule ligand nanopatterning platform, we find that cardiomyocytes spreading on laminin binding integrins require more concentrated integrin clusters to initiate spreading, compared to RGD binding integrins. Our data further indicates differences in the requirements for local (nanometre scale) vs global (micrometre scale) ligand densities and that these differences are driven by the respective stability of the adhesions to existing levels of cytoskeletal contractility.

While the functional implications of differential integrin expression in cardiomyocytes are poorly understood, we can look to other cell types that modulate integrin expression profiles to perform specific functions (e.g. migration, adhesion formation). The different integrins possess unique ECM-bond dynamics (e.g. binding/unbinding rates & catch/slip bond behaviour), which are highly sensitive to force (i.e. contractility and substrate rigidity) and are further characterised by their nanoscale organisation and clustering properties[15, 26].

Especially, αvβ3 expressing cells required an inter-ligand distance between individual RGD molecules of 60nm or less, disregarding the global concentration, in order to form stable adhesions[15, 27] [27-31]. Moreover, a study employing peptidomimetic ligands specific for αvβ3 and α5β1 integrins found similar spreading for both integrin types with 60nm inter-ligand spacing but detected further spreading of α5β1 integrins at even smaller ligand spacings[15].

Little was known about the clustering behaviours of fibronectin and laminin integrin subtypes in cardiomyocytes, especially α5β1 and α7β1 [32, 33]. Here, we observed the same 60 nm threshold for RGD binding integrins. However, unlike observations in fibroblasts with larger levels of αvβ3 integrins and differences in β1 integrin and talin isoforms[21], we found a strong contribution of global in addition to local ligand densities and in fact cardiomyocytes were spread out to same extents on similar global concentrations of RGD peptides, independent of the spacing of the ligands on each cluster or the distance between the clusters. Intriguingly, for cardiomyocytes laminin binding integrins required much smaller spacings of ∼30 nm (or 12 peptides per cluster) to initiate spreading, compared to the fibronectin binding integrins. Moreover, there seemed to be a stricter requirement for the local, in addition to global ligand concentrations. This suggested a difference in the ability to form adhesions over multiple clusters and stably distribute the adhesion forces over larger distances between the clusters (see model in Figure 7). In other words, for RGD binding integrins, a 200nm inter-cluster spacing can be bridged with only 6 ligands per cluster; however at least 12 ligands per cluster are required in order to bridge the 300nm distance. A stronger requirement for inter-ligand vs inter-cluster spacing for IKVAV binding integrins in turn, agrees with an overall lower adhesion force and stability and hence an inability to effectively bridge clusters even at 200nm distance.

Integrin clustering is a critical part of adhesion formation and responds to molecular tension. Integrins demonstrate typical catch bond behaviour and extend the bond lifetimes to their ligand under the application of force[6]. Additionally, the integrin binding protein talin is stretched in a force dependent way leading to the opening of cryptic vinculin binding sites[19]. Vinculin then provides additional links between talin and actin to reinforce the adhesion[6]. However, if the forces, or loading rates are too high, then bonds will break before the adhesion can form. Integrin clustering directly modulates the forces experienced by components of the integrin adhesion; a greater number of integrins enables distribution of traction forces among more proteins, resulting in less force being sensed by individual integrins and adaptors[26].

Pharmacological inhibition of cytoskeletal contractility resulted in an increase in NRC spreading on IKVAV, further suggesting that laminin binding integrin adhesions (at current ligand densities) may be less stable under high forces. This would also agree with observations of decreased cytoskeletal content, traction forces and spreading in cells cultured on laminin.

Together, these results suggest that fibronectin adhesions mediate stronger spreading, contractility, and integrin adaptor enrichment and enhanced stability of individual adhesions. Furthermore, this finding suggests that a transition in integrin expression to favour fibronectin binding subtypes (especially α5β1) may also alter integrin signalling and mechanotransduction pathways [34, 35].

To specifically analyse the integrin-ligand interactions, we used here micro- and nanopatterned 2D surfaces. The use of 2D surfaces comes with the limitation that upon plating, cardiomyocytes remodel to form peripheral focal adhesion structures in addition to costameric structures at the level of the z-disc (e.g. visible by a striated vinculin staining)[5]. Nevertheless, the presence of fibronectin and laminin binding integrins make neonatal rat cardiomyocytes an excellent model to study the nanoscale adhesion properties. However, future work will need to analyse the contributions of the different β1 integrin and talin isoforms, which differ in their affinities to each other[16].

In summary, the current results provide novel insights into the cardiomyocyte integrin adhesions during postnatal heart development or in heart disease, where we find that the switch in cardiomyocyte integrin expression has important implications for integrin signalling and mechanotransduction.

## Methods

### Cells

Neonatal rat cardiomyocytes were prepared as described previously[5]. Hearts were dissected into ice-cold ADS buffer (116mM NaCl, 20mM Hepes, 0.8mM NaH_2_PO_4_, 5.6mM Glucose, 5.4 mM KCL, 0.8 mM MgSO_4_). After hearts settled down to the bottom, hearts were washed once with ADS buffer. ADS buffer was then removed, and hearts were incubated with 5ml enzyme solution in ADS (ES, 246U Collagenase and 0.6 mg Pancreatin / ml), for 5 min, at 37C under vigorous shaking. Supernatant was discarded. This step was followed by 5-6 digests, until hearts were completely digested. Each time 5ml fresh ES was added to the hearts and incubated 15min at 37C, under shaking. Hearts were pipetted up and down 30 times using a pasteur pipette. After settling down, supernatant was transferred into plating medium (65% DMEM, 17% M199, 10% Horse Serum, 5% FCS, 2% Glutamax, 1% Penecillin/Streptamycin (P/S)). Two digests each were combined in one tube with 20ml plating medium, then cleared through a 100*μ*m cell strainer and spun down at 1200rpm for 5min at RT, before resuspended in 10ml plating medium. Cells were pooled together and pre-plated for 90min to enrich the cardiomyocytes. Cardiomyocytes were then plated onto the respective substrates as indicated in the text or figures. Medium was changed the next day to maintenance medium (77% DMEM, 18% M199, 2% Horse Serum, 2% Glutamax, 1% P/S), or serum starvation medium (as above, but excluding the Horse Serum).

The following drugs were applied for 2h in serum free medium: Y27632 (10 *μ*m; Tocris); Blebbistatin (5 *μ*m; Calbiochem); Calyculin A (100 nM; Sigma). Adult rat cardiomyocytes were isolated by Langendorff perfusion of hearts as described previously[17]. Adult and newborn rats were sacrificed in accordance with the Schedule 1 to the Animals (Scientific Procedures) Act 1986.

### Immunostainings

For immunostaining, cells were fixed with 4% PFA for 10 minutes, permeabilized with 0.2% triton X-100 in PBS for 5 minutes, blocked with 5% BSA in PBS for 1h and stained in the antibody solutions in immunostaining buffer (20 mM Tris, 155 mM NaCl, 2 mM EGTA, 2 mM MgCl2, 1% BSA at pH 7.4) [17]. Cells were washed three times for 10 minutes with PBS after each step and mounted in MOWIOL 4-88 (0.1g/ml in Glycerol/Water/Tris (0.2M, pH8.0) at a ratio of 1/1/2) containing a final concentration of 4% n-propyl gallate. Live cell imaging and imaging of multiwell plates was performed on an inverted Nikon Eclipse Ti-E microscope with a Nikon DS-Qi2 sCMOS camera and equipped with a Solent Scientific chamber with temperature and CO2 control. Confocal microscopy was performed on a Nikon A1R+ inverted microscope with GaAsP Detectors. TIRF imaging and bleaching curves were recorded on a Zeiss LSM710 Elyra Microscope.

### PDMS substrates and nanopillars

PDMS pillar (500 nm diameter, 1.7 *μ*m height, 1*μ*m centre-to-centre) substrates were prepared by soft lithography from silicon masters as described previously[36]. For fluorescent labelled pillars, CdSeS/ZnS alloyed quantum dots (490nm, Sigma) were spun first on the master 30s at 10,000rpm with a 150i spin processor (SPS), before the addition of PDMS. PDMS (Sylgard 184, Dow Corning) was mixed thoroughly with its curing agent (10:1), degassed, poured over the silicon master, placed upside-down on a plasma-treated coverslip-dish (Mattek), or coverslip 4-well dishes (Ibidi) and cured at 80C for 2 h. The mould was then removed and the pillars were incubated with fibronectin for 1h at 37C.

Flat PDMS substrates were prepared by spin-coating Sylgard 184, Sylard 527 or mixtures at the Ratios of 1:5, 1:10 and 1:20 with a 150i spin processor (SPS), onto coverslips for western blotting samples, or microscope slides for placing into multiwell plates (Grace Biolabs). Before spin coating, Sylgard 527 was pre-cured at 70C for 30 minutes with intermitted mixing to achieve a comparable viscosity to the Sylgard 184 mixture.

### Micropatterning

Microcontact printing was performed as described elsewhere[18]. Briefly, hPDMS stamps were cast on E-beam-lithographed PMMA wafers. 20 μg/ml ligand proteins in PBS were deposited onto the stamps for 40 min. Micro-patterns were printed by stamping on plasma-treated glass for 1 min. Fibronectin/ laminin micro-grids were printed in two steps: (1) fluorescent fibronectin deposition followed by (2) printing of transverse fluorescent laminin lines. Cells were seeded after washing and blocking with 5%BSA.

### DNA Origami

DNA origami was assembled as described previously[37]. Briefly, M13mp18 (5 nM) and staple strands (50nM) were combined in 50μL of TAE buffer with 12.5 mM Mg^2+^ (DNA sequences can be obtained at [38]). M13mp18 is a bacteria phage vector strand with 7249 bases long. An appropriate quantity of ions, such as magnesium here, or sodium, are required to efficient DNA hybridization. This acts to equilibrate electrostatic repulsion between highly negatively charged DNAs molecules. An amount of 12.5 mM Mg^2+^ sufficient to achieve a high yield of DNA origami and limit any aggregation effects. DNA origami are synthesised by annealing from an initial temperature of 94 °C to completely melt all dsDNA. Temperature step-controlled annealing was carried out in a PCR machine. Samples were cooled from 94 to 65 °Cat a rate of ∼0.3 °C per minute. A cooling rate of 0.1 °C per minute is employed from 65 °C to room temperature. The self-assembled DNA origami were then purified using Millipore Amicon Ultra 100 kDa spin columns in a centrifuge at 2000 rcf for 6 min, three times, to remove excess staple strands. DNA origami were adjusted to a concentration ∼20 nM and stored in Lo-Bind Eppendorf tubes at 4 °C. A NanoDrop spectrophotometer is used to detect the approximate concentration of DNA origami products based on the constant of a molecular weight of 330 g/mol per base and an extinction coefficient = 33 mg/mL for A260 = the actual result is close to the estimated numbers. To ensure efficient assembly and labelling of DNA origami with peptide conjugates, unmodified staple strands and amino anchors were added at a 5× excess to the M13mp18 backbone and peptide conjugates were added at a 10× excess.

RGD and IKVAV peptides were conjugated to ssDNA through UV mediated thiol-ene reaction, via a sulfhydryl group on the peptides and a thiol group on acrydite modified ssDNA (Integrated DNA Technologies). Peptides were diluted to 100*μ*m in H_2_O containing 100× TCEP pH 7.0, to reduce any disulfide bonds and present the cysteine groups for conjugation. Peptides were then mixed with the acrydite modified ssDNA at a final concentration of 20 *μ*m and 200 *μ*m, respectively. Reactions were carried out in 120mM Tris buffer with 11 *μ*m photoinitiator (2-hydroxy-4′-(2-hydroxyethoxy)-2-methylpropiophenone). Samples were exposed to 260 nm UV light for 1 hour and the conjugates were purified by RP-HPLC and freeze drying, as described above.

DNA origami were characterised by AFM (Bruker, Dimension Icon) as described previously**[37]**.

### Peptide and Amino Modifications to DNA Origami

The pointed triangle design was selected due to its lower tendency to aggregate and to exhibit blunt end stacking effects[39]. Eighteen peptide and 15 amino anchor modifications were made to the triangle origami design. For the peptide modifications, the original strands that hybridise to the m13m18 backbone were extended with the same sequence (AGTTGTGGATCCTACT). This extended sequence was complementary to the acrydite peptide linker when was conjugated to the peptide. Strands selected for the amino anchors were modified with a poly-T extension, complementary for poly-A 6 carbon amino anchor. All modifications were reported previously[37]. During synthesis of the DNA origami, peptide and amino modification were added at a 20x excess to the concentration of the available modification sites.

### Nanopatterning

Nanopatterns were produced by direct write e-beam lithography (EBL) on coverslips. Coverslips were solvent cleaned in an ultrasonic bath (acetone followed by methanol, IPA, and water) and dehydrated in a 180C oven overnight. An oxygen plasma treatment for 60s at 100W power prepared the surface for silane deposition, which was done from HMDS vapour in a closed container at 150 C. A 950,000 g/mol poly(methyl methacrylate) (PMMA) film was spin coated on the coverslips at 5000 rpm, followed by a short solvent bake on a 180C hotplate for 30s. A 10nm aluminium charge conduction layer was evaporated on the coverslips, and they were mounted on a 4-inch silicon wafer for EBL processing using crystalbond 555 adhesive. EBL exposure was carried out as described previously[40] to define arrays of 200nm and 300nm pitch holes, diameter 150nm in the PMMA layer. After exposure, the aluminium charge conduction layer was removed in a 2.6% TMAH solution (MF-CD26) for 30s followed by water rinse.

For patterning of DNA origami on e-beam patterned substrates, patterned substrates were first developed in a 2:1 solution of propanol-2:methyl isobutyl ketone for 60 seconds at 23°C, followed by a 30 seconds wash in 100% propanol-2. DNA origami binding sites were then etched using oxygen plasma treatment (100W, 70 seconds, room air) followed by silanisation with 0.1% CTES in Tris buffer (5 mM, pH 8.0). The PMMA resist was then removed by immersion in NMP at 50 °C and sonicated or 10 minutes. DNA Origami were incubated for 1 hour at room temperature at a concentration of 1 nM in Tris buffer (5 mM, pH 8.3, 50 mM MgCl_2_, see supplementary methods for more information on placement conditions). Due to the hydrophobic nature of the HMDS layer and the size of the patterned area, 100 μL of origami solution was required to cover the surface sufficiently. Samples were then washed with MOPS buffer (10 mM, pH 8.1, 50 mM MgCl_2_) to remove primary amines, followed by a wash with MOPS buffer (10 mM, pH 8.1, 50 mM MgCl_2_) containing 50 mM EDC and 25 mM sulfo-NHS. Finally, samples were then washed with the MOPS buffer without MgCl_2_and rinsed with DPBS containing 125 mM NaCl. Before AFM characterisation, samples were placed into deionised H_2_O.

### Image Analysis

Pillar displacements were analysed with imageJ, using the NanoTracking plugin. An image of the pillars after removal of the cells with 10x trypsin was taken as reference for the non-displaced pillars. For contraction analysis, pillar displacements from spontaneous contracting cardiomyocytes were measured for the whole movie, using Matlab. From the data, the maximum displacement (systole) was compared to the subsequent minimum and to the non-displaced pillars. Noise levels were measured from pillars outside the cell and estimated around 30nm (consistent with our previous work)[5]. Therefore, all pillars that were displaced above 30nm during the movie were considered for the analysis. Statistics were calculated between cells. Cell area, cell morphology and staining intensity were analysed with cell profiler.

### Quantification and statistical analysis

Data sets were tested for normal distribution using the Shapiro–Wilk test. All box plots are displayed as median (central line), upper and lower quartile (box), +/- 1.5 x inter quartile range (whiskers). All n-numbers and statistical tests are indicated in the figure legends. All statistic tests were performed with Graphpad Prism.

## Acknowledgments

We would like to acknowledge the James Watt Nanofabrication Centre and its staff at the University of Glasgow for the nanofabrication work. WH is recipient of a BBSRC LIDO Studentship. DH was supported by the China Scholarship Council. TI was supported by a British Heart Foundation Intermediate Basic Science Research Fellowship (FS/14/30/30917), BHF project grant (PG/20/6/34835) and a BBSRC new investigator award (BB/S001123/1). NG also acknowledges ERC funding through FAKIR 648892 Consolidator Award.

