## Supplementary Information for "Regulation of Cardiomyocyte Adhesion and Mechanosignalling Through Distinct Nanoscale Behaviour of Integrin Ligands Mimicking Healthy or Fibrotic ECM"

**Supplementary Figures:**

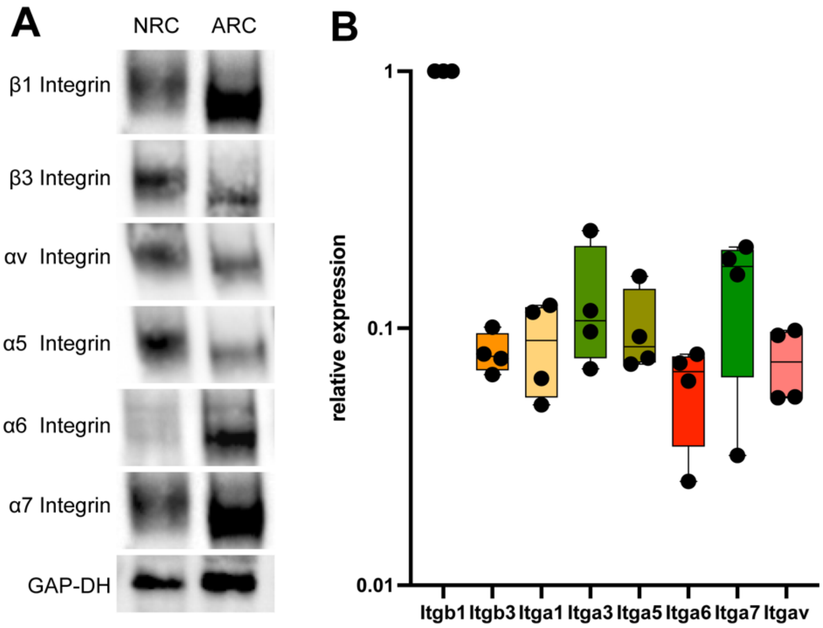


**Supplementary Figure S1:** Fibronectin and Laminin binding integrins are expressed in neonatal rat cardiomyocytes. A) Western blot comparison of neonatal (NRC) and adult ventricular cardiomyocytes (ARC). Fibronectin binding α5 and αv integrins are more strongly expressed while β1, α6 and α7 are expressed less in neonatal rat cardiomyocytes (NRC), compared to adult cardiomyocytes (ARC). **B)** Meta-study of integrin expression levels in neonatal rat cardiomyocytes, displayed as relative expression in comparison to β1 integrin. Both fibronectin and laminin binding integrins are expressed at comparable levels in neonatal rat cardiomyocytes. Integrin expression levels were collated from total RNA expression data from untreated control neonatal rat cardiomyocytes from the following data sets (displayed as average per data set): GDS3763 (3 control samples)[1], GDS3117 (2 samples)[2], E-MEXP-2516 (5 samples)[3] and E-MEXP-2527 (4 samples)[4].

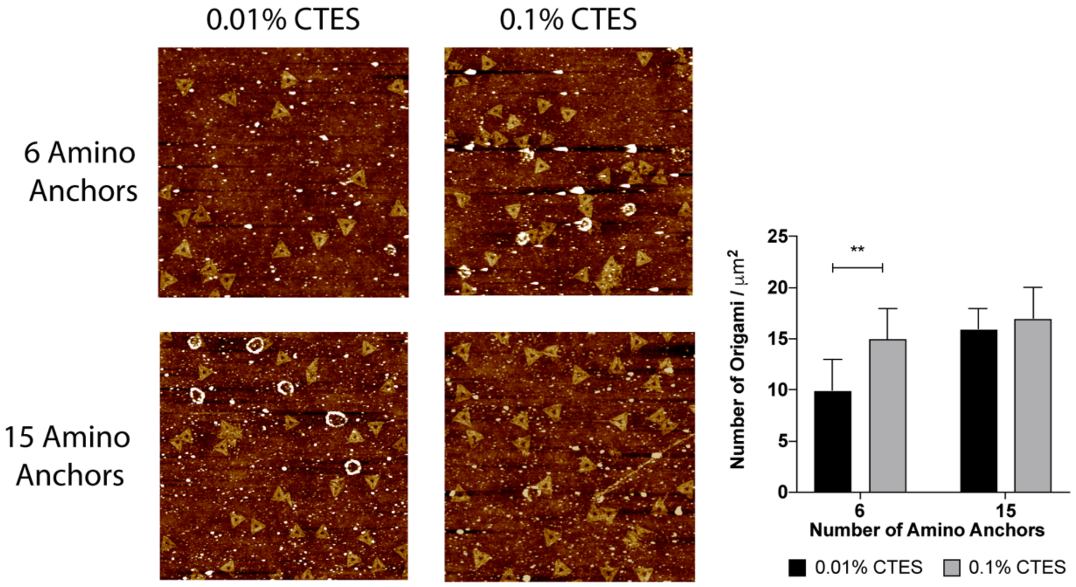


**Supplementary Figure S2:** Optimisation of surface silanisation results in higher cross-linking yield, independent of the number of amino anchors. Glass surfaces were silanised with 0.01% CTES or 0.1% CTES for 30 minutes. AFM was then used to image the DNA DNA origami presenting 6 or 15 amino anchors which had been cross-linked to the surface. The quantification displays mean Å} SD DNA origami density per μm2 over three independent repeats. Student's t-test, ** = p<0.01.

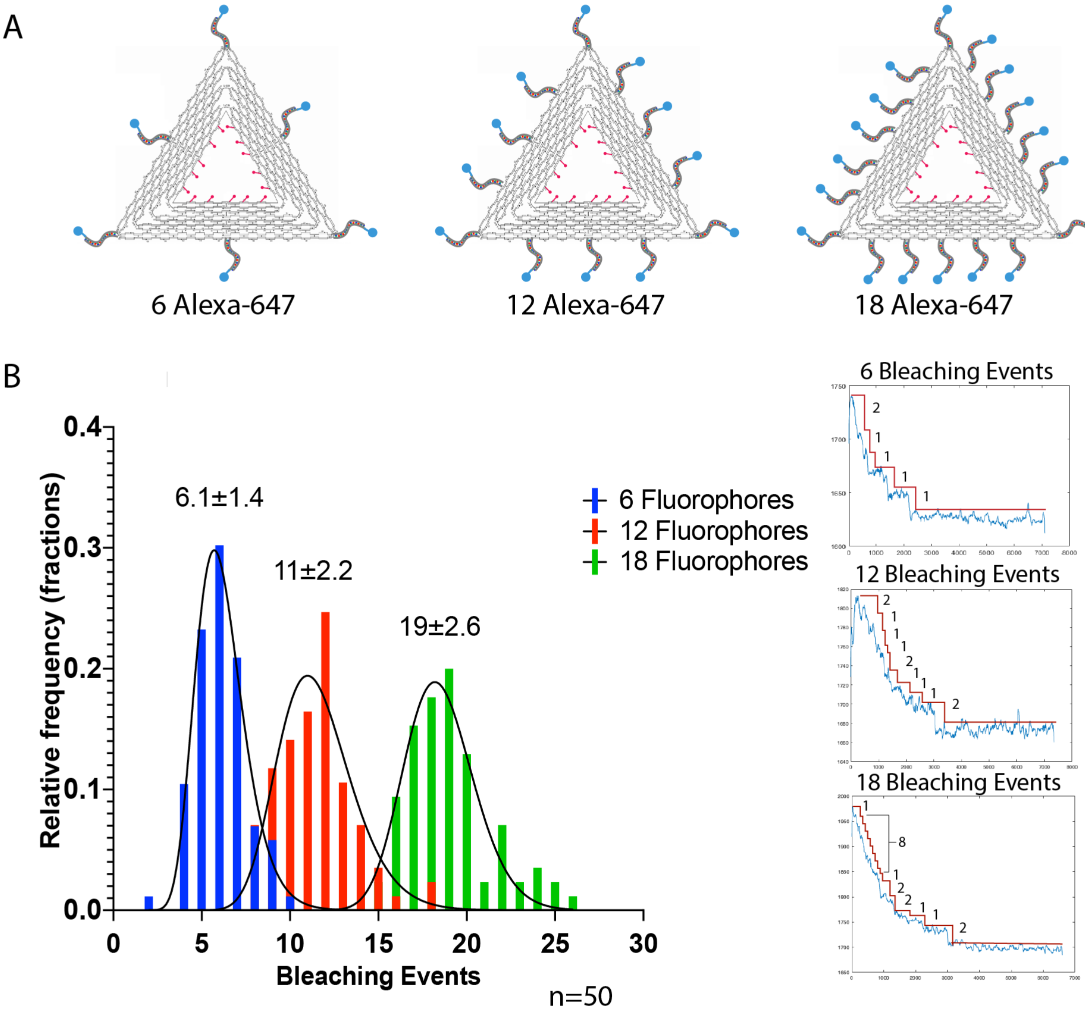


**Supplementary Figure S3:** Validation of the peptide attachment approach using photobleaching of DNA origami labelled with Alexa-647 fluorophores. A) DNA origami were labelled with 6, 12 or 18 Alexa-647 fluorophores and cast onto glass bottomed dishes, loaded onto a confocal microscope and photobleached. B) A histogram of the number of bleaching events revealed an average of 6, 11 and 19 fluorophores in the 6, 12 and 18 fluorophore conditions. N = 50 analysed cluster per condition. Frame rate = 140 fps.

| Configuration | Ligands per DNA structure | Inter-peptide distance | DNA structure spacing | Global ligand Density |
|---|---|---|---|---|
| 6/Random | 6 | ~60 | Variable | ~60 |
| 18/Random | 18 | ~20 | Variable | ~180 |
| 6/300 | 6 | ~60 | 300 | 69 |
| 6/200 | 6 | ~60 | 200 | 150 |
| 12/300 | 12 | ~30 | 300 | 139 |
| 12/200 | 12 | ~30 | 200 | 300 |
| 18/300 | 18 | ~20 | 300 | 208 |
| 18/200 | 18 | ~20 | 200 | 450 |

**Supplementary Table 1:** Overview over bionanoarray configurations
